# Spatial signature of white matter hyperintensities in stroke patients

**DOI:** 10.1101/457374

**Authors:** Markus D. Schirmer, Anne-Katrin Giese, Panagiotis Fotiadis, Mark R. Etherton, Lisa Cloonan, Anand Viswanathan, Steven M. Greenberg, Ona Wu, Natalia S. Rost

## Abstract

**Purpose:** White matter hyperintensity (WMH) is a common phenotype across a variety of neurological diseases, particularly prevalent in stroke patients; however, vascular territory dependent variation in WMH burden has not yet been identified. Here, we sought to investigate the spatial specificity of WMH burden in patients with acute ischemic stroke (AIS).

**Materials and Methods:** We created a novel age-appropriate high-resolution brain template and anatomically delineated the cerebral vascular territories. We used WMH masks derived from the clinical T2 FLAIR MRI scans and spatial normalization of the template to discriminate between WMH volume within each subject’s anterior cerebral artery (ACA), middle cerebral artery (MCA), and posterior cerebral artery (PCA) territories. Linear regression modeling including age, sex, common vascular risk factors, and TOAST stroke subtypes was used to assess for spatial specificity of WMH volume (WMHv) in a cohort of 882 AIS patients.

**Results:** Mean age of this cohort was 65.23±14.79 years, 61.7% were male, 63.6% were hypertensive, 35.8% never smoked. Mean WMHv was 11.58c +/-13.49 cc. There were significant differences in territory-specific, relative to global, WMH burden. In contrast to PCA territory, age (0.018±0.002, p<0.001) and small-vessel stroke subtype (0.212±0.098, p<0.001) were associated with relative increase of WMH burden within the anterior (ACA and MCA) territories, whereas male sex (−0.275±0.067, p<0.001) was associated with a relative decrease in WMHv.

**Conclusions:** Our data establish the spatial specificity of WMH distribution in relation to vascular territory and risk factor exposure in AIS patients and offer new insights into the underlying pathology.

## 1 Introduction

White matter hyperintensity (WMH) is an important and widely studied radiographic phenotype (Debette and Markus, 2010). Even though WMH burden has been linked to both incidence and outcomes of stroke (Kuller et al., 2004), spatial specificity of WMH has not been fully explored. WMH is commonly assessed on axial T2 Fluid Attenuated Inverse Recovery (FLAIR) magnetic resonance (MR) images and total WMH burden can be summarized as WMH volume (WMHv) (Debette and Markus, 2010; Giese et al., 2017; Kuller et al., 2004). Importantly, it has been shown, that WMHv presents an accurate and uniform way of quantifying this phenotype in clinical populations, such as acute ischemic stroke (AIS) patients (Zhang et al., 2015). However, summarizing the burden on such a high level, i.e. total WMHv, eliminates potential spatial specificity, which in turn may provide insight with regard to underlying vascular pathology and disease progression.

While other studies have explored spatial WMH patterns in diseased and healthy populations (Griffanti et al., 2017; Holland et al., 2008; Kim et al., 2008), prior studies exploring the extent of WMH on a voxel-based level in AIS patients with respect to varying risk factors, such as hypertension (Yoshita et al., 2006), were limited due to high dimensionality. For WMH, as a prototypical small vessel disease (Rost et al., 2010), a spatial differentiation may be based on supplying arteries, as previously established for stroke lesions (Ng et al., 2007). In the human brain, we distinguish between the anterior and posterior circulation. The anterior supratentorial circulation is comprised of the anterior (ACA) and middle cerebral artery (MCA), while the posterior cerebral artery (PCA) supplies the posterior supratentorial vascular area. Infratentorial structures receive their blood supply through the posterior circulation. Moreover, large-scale analysis of WMH burden for each vascular territory is limited, as manual assessment and differentiation of individual vascular territories is time consuming and results in inter-rater variations.

In general, medical image analysis is prone to inter-subject variability. It is common practice to reduce this variability by normalizing images to a common coordinate system (Mandal et al., 2012). In addition to allowing reproducible assessment of disease phenotypes across subjects with respect to their spatial location, it furthermore allows comparison across studies. Moreover, computational costs can be reduced by utilizing prior information such as brain or tissue segmentation. However, for WMH analysis in AIS patients, an age-appropriate T2 FLAIR template for spatial normalization with outlined vascular territories is currently missing.

Here, we examined the distribution of WMH, differentiated by supratentorial cerebral vascular territories in clinically acquired axial T2 FLAIR images of patients with AIS. In order to assess spatial distribution, we first created an age appropriate brain template, including T1, T2 and 3D-FLAIR sequences based on high-resolution MR images, and delineate all 5 bi-lateral vascular territories. We utilized these vascular territories, by spatially normalizing each subject’s axial T2 FLAIR image to the 3D-FLAIR template. Finally, we investigated WMHv for each of the supratentorial vascular territories with respect to the common clinical WMH risk factors, while accounting for territory size, in a multivariate analysis.

## 2 Materials and methods

### 2.1 Study design and patient population

Patients were enrolled as part of the Genes Associated with Stroke Risk and Outcomes Study (GASROS) between 2003 and 2011 (Zhang et al., 2015). Patients presenting to the Massachusetts General Hospital Emergency Department (ED) within 12 hours of AIS symptom onset and who were >18 years of age, were eligible to enroll. Each patient was evaluated by a vascular neurologist and clinical variables, including age, sex, common vascular risk factors (history of hypertension (HTN), diabetes mellitus (DM2), hyperlipidemia (HLD), tobacco smoking) and TOAST stroke subtypes (CE: cardioembolic, LA: large-artery atherosclerosis, SV: small-vessel occlusion, Other: other determined etiology; (Adams et al., 1993)) were recorded. Each patient underwent standard clinical imaging protocol within 48 hours of admission, including axial T2 FLAIR imaging (TR 5000ms, minimum TE of 62 to 116ms, TI 2200ms, FOV 220-240mm). This study was carried out in accordance with the recommendations of Partners Institutional Review Board with written informed consent from all subjects. All subjects gave written informed consent in accordance with the Declaration of Helsinki. The protocol was approved by the Partners Institutional Review Board.

We identified 882 subjects with manual WMH outlines (Zhang et al., 2015) and most complete phenotypic information available for this analysis (Table 1). Manual WMH outlines were performed using MRIcro software for computer-assisted determination of WMHv (Rost et al., 2010). Maps were created using axial T2 FLAIR sequences, based on a previously published semi-automated method with high inter-rater reliability (Chen et al., 2006). Each subject’s DWI sequence was utilized to exclude acute ischemia, edema, and chronic infarcts. Additionally, out of the 882 subjects with confirmed DWI lesions, 586 subjects had lesions manually outlined, using a semi-automated algorithm (Mocking et al., 2011), by reader blinded to the admission stroke severity and 90-day outcome, measured by the modified Rankin Scale.

**Table 1:**
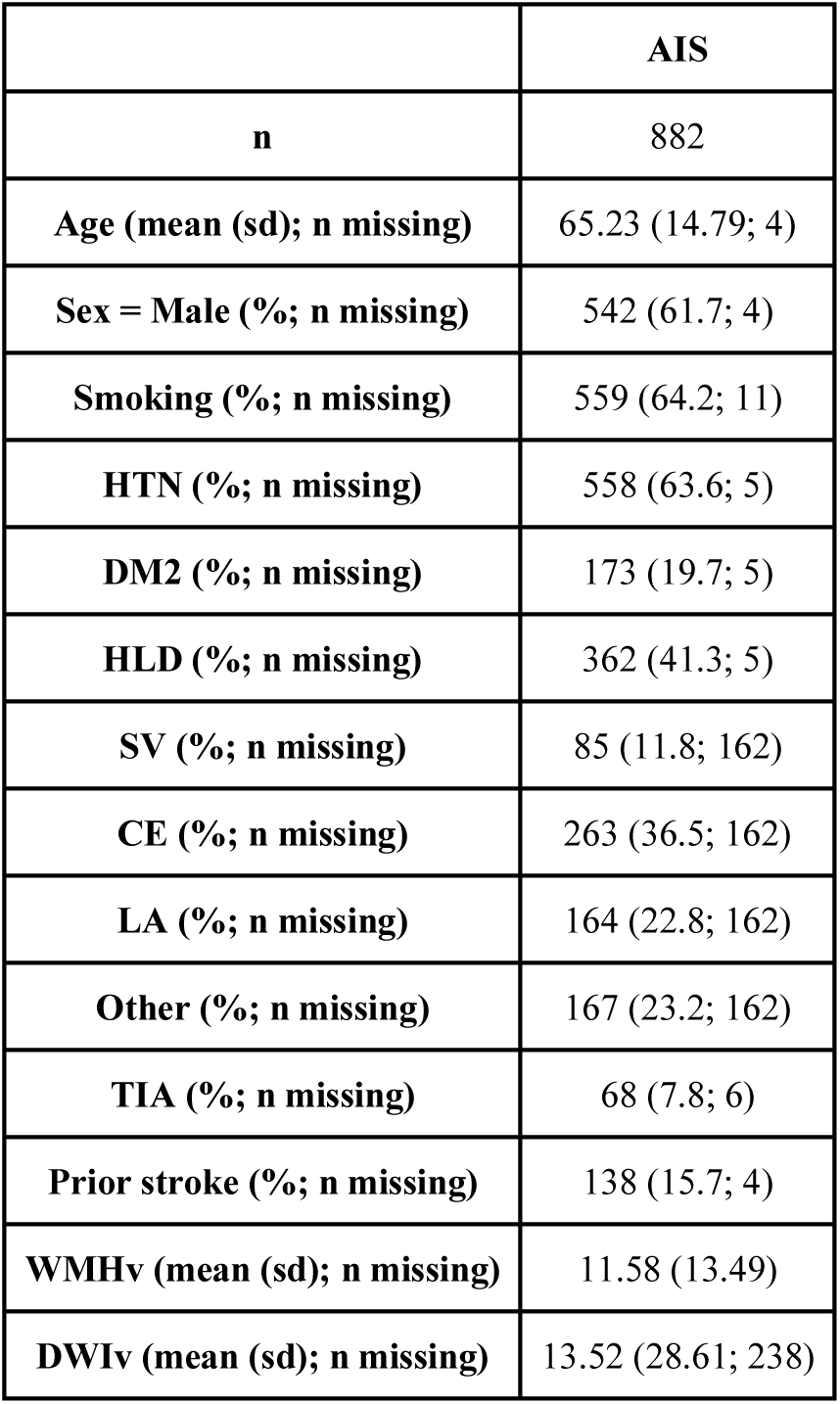
Study cohort characterization. Smoking is assessed based on part or current history of smoking (ever/never).

In addition to the AIS cohort, 16 subjects were recruited between 2016 and 2017 at Massachusetts General Hospital, to generate a high-resolution template. The template cohort selection aimed to match characteristics (age and sex) of the elderly adults presenting to the ED with pre-existing cerebrovascular pathology and incident AIS. Twelve stroke-free, non-demented patients with the sporadic form of cerebral amyloid angiopathy (CAA), a known cerebral small vessel disorder and similarly-aged healthy controls (n=4) underwent high-resolution MRI as part of a separate study at our hospital (mean (standard deviation) age: 69.6 (8.2); 62.5% male). MR scans were manually assessed to exonerate any gross pathology, such as hemorrhage or silent brain infarcts.

High-resolution structural brain MRI sequences were acquired with a Siemens Magnetom Prisma 3T scanner (using a 32-channel head coil). The standardized protocol included a Multiecho T1-weighted (voxel size: 1×1×1 mm^3^; Repetition Time [TR]: 2510 ms), a 3D-FLAIR (voxel size: 0.9×0.9×0.9 mm^3^; TR: 5000 ms; TE: 356 ms), and a T2-weighted Turbo Spin Echo (voxel size: 0.5×0.5×2.0 mm^3^; TR: 7500 ms; TE: 84 ms) sequence.

### 2.2 Template creation

We employed Advanced Normalization Tools (ANTs) for image processing, a well-established tool for image registration and template creation (Avants et al., 2010, 2011). Utilizing the 16 high-resolution images, we created a brain template based on multimodal information using T1, T2 and 3D-FLAIR sequences. After template creation was completed, and due to the relatively low number of subjects, we smoothed the resulting templates (FSL; Gaussian smoothing, sigma = 1). Finally, we registered the resulting templates into MNI space using ANTs (Avants et al., 2011).

Additionally, manual brain extraction was performed on all subjects based on their 3D-FLAIR sequences. Binary manual masks were warped into template space and averaged. For each voxel, majority voting was performed and an average brain mask was generated based on the voxels where more than 50% of the subjects agreed, providing an initial brain mask, which was manually assessed and corrected on a per-slice basis where necessary.

Vascular territories were outlined on the right hemisphere in the T1-weighted atlas image by an expert neurologist (A.K.G.). Vascular territories included anatomically validated ACA, MCA, and PCA territories supratentorially. We also created the anterior territory (ANT), a combined ACA and MCA territory named after the “anterior” circulation versus “posterior” supratentorial circulation supplied by the PCA. The outline was then mirrored onto the left hemisphere and subsequently combined to create a full-brain vascular territory map. This map was manually assessed and corrected where necessary. Additionally, WMHv and ventricular size were manually determined on the atlas, to ensure that the template is representative of subjects in this age category.

### 2.3 Neuroimage analysis of WMH burden

Each axial T2 FLAIR image was first skull-stripped using an in-house skull-stripper for clinical quality scans (Schirmer et al., 2017) and then non-linearly registered to the high-resolution 3D-FLAIR template using ANTs (Avants et al., 2011). Subsequently, the vascular territory map was warped into subject space, allowing us to calculate both the volume of each vascular territory in subject space, as well as the WMH burden for each territory. Finally, we normalized each territorial WMHv by the total WMHv, to describe a relative burden per territory (WMHv_rel_).

Each subject’s vascular territory map was manually assessed for gross registration errors, such as midline shift. Furthermore, we identified potential outliers using the modified z-score (Iglewicz and Hoaglin, 1993) for each set of territorial volumes of the cohort. Subjects with registration errors and those deemed outliers were removed from analysis.

Additionally, we registered all 586 subjects non-linearly with manual DWI lesion outlines to the 3D-FLAIR template. This allowed us, with the addition of WMH outlines being mapped to the same template, to create incidence maps of both the chronic WMH disease burden, as well as the acute lesion locations.

### 2.4 Statistical analysis

First, we transformed the relative WMH disease burden in each territory using a logit transformation, to avoid potential issues due to the response variables in the models being bound between 0 and 1. We then assessed WMH burden for each territory based on a univariate analysis, where each phenotype was used in a linear model (continuous) or Mann-Whitney U test (categorical) for each of the transformed relative WMH burden. This was followed by a multivariate analysis, where all factors were included in the model, given by

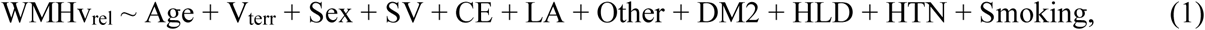

including age, volume of the territory in subject space (V_terr_), sex, TOAST subtypes (yes/no) (including SV, LA, CE, Other), DM2, HLD, HTN, and smoking (ever/never) status. The undetermined TOAST subgroup was not investigated in this analysis. To reduce the statistical burden of utilizing 11 independent variables in the model, we used a backward selection approach to simultaneously refine all 4 territorial models, i.e. MCA, ACA, PCA, and ANT, by removing the variable with the highest p-value above 0.1 iteratively, where p-values were summarized as the minimum p-value for the variable across the models.

As a comparison, we also investigate the efficacy of this model, compared to a baseline model, where the WMHv_rel_ is solely explained by V_terr_, given by

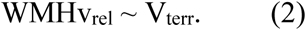

Both models are compared using a *χ*^2^ test.

Statistical analysis was conducted using the computing environment R (R Core Team, 2013).

### 2.5 Data availability

The utilized template will be made available upon acceptance to facilitate reproducibility of our findings. The authors agree to make available to any researcher the data, methods used in the analysis, and materials used to conduct the research for the express purposes of reproducing the results and with the explicit permission for data sharing by the local institutional review board.

## 3 Results

### 3.1 Template and territorial map

Utilizing ANTs, we created a multimodal template from the 16 subjects with high-resolution MR images. Figure 1 shows axial slices of the T1, T2 and 3D-FLAIR template, after registration to MNI space. Additionally, it shows the vascular territory map, which was delineated on the template’s T1 sequence. Within this template, WMH and ventricular volume were manually assessed, with 13.2cc and 34.4cc, respectively.

**Figure 1:**
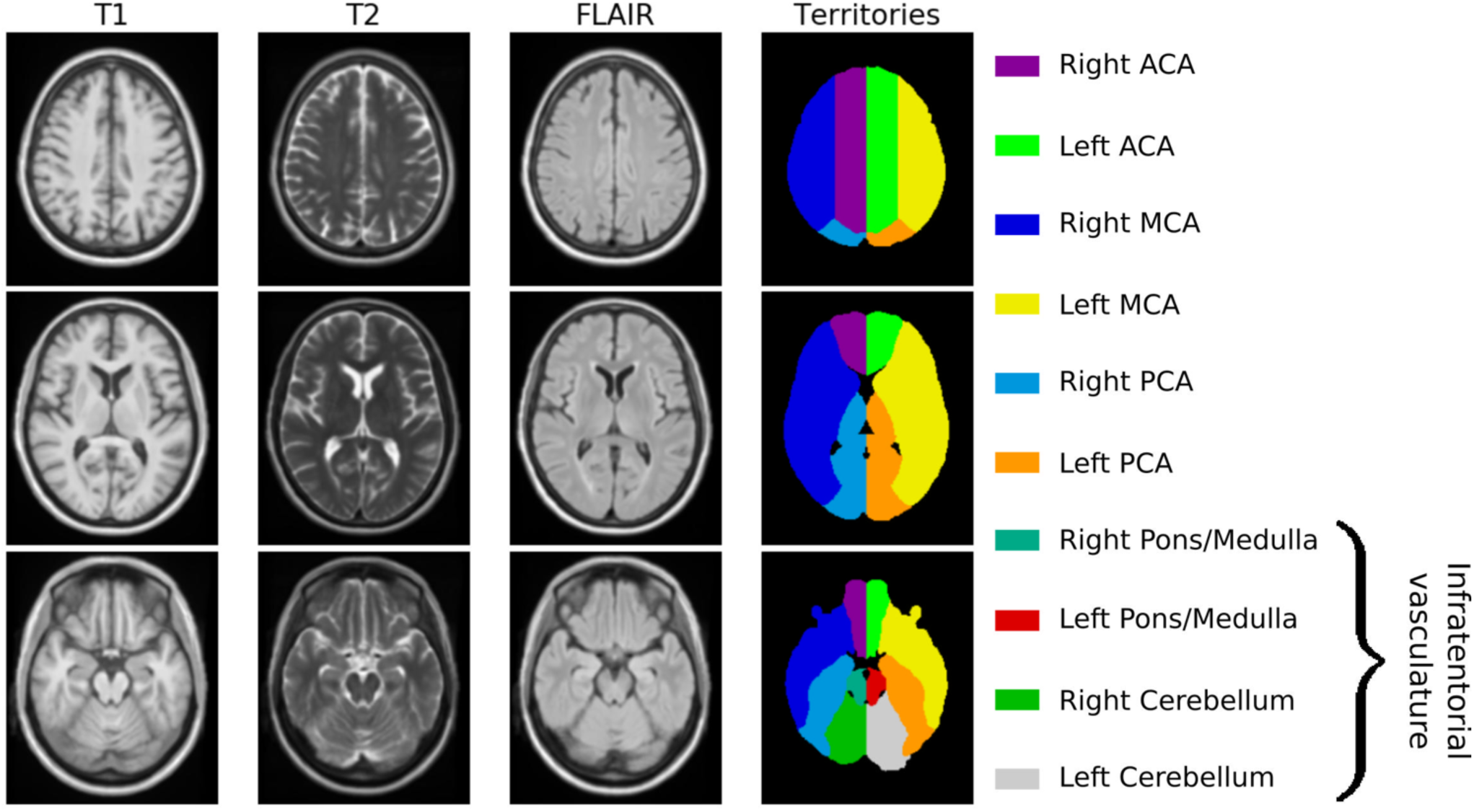
Age-appropriate template, including T1, T2 and 3D-FLAIR sequences for image registration, as well as vascular territory map.

### 3.2 Neuroimage analysis of WMH burden

#### Final cohort and territorial WMH burden characterization

Each subject is non-linearly registered to the 3D-FLAIR template and the vascular territory map transformed into low-resolution subject space. We assessed each subject for registration errors, leading to 17 subjects being excluded. Utilizing the modified z-score for each territory’s volumes did not lead to further exclusions. Registering the DWI volumes to the 3D-FLAIR template did not lead to any of the 586 subjects being excluded. Figure 2 shows the spatial distribution of WMH burden and acute lesion location within the template.

**Figure 2:**
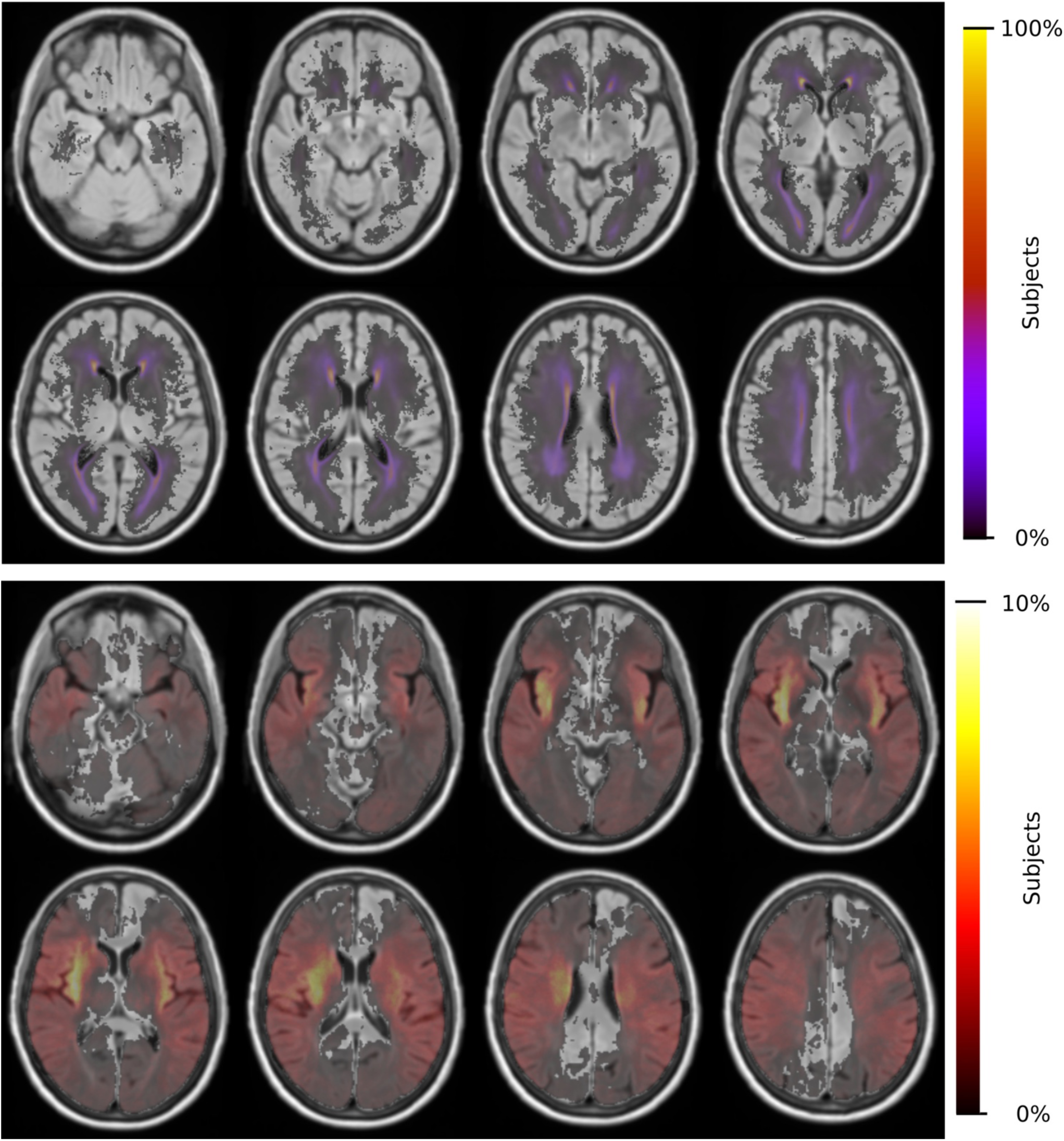
Distributions of WMH burden (top) and acute lesion location (bottom) for 865 patients and 586 patients, respectively.

For the remaining 865 subjects with WMH outlines, ACA, MCA and PCA volumes (mean ± standard deviation) were estimated to be 334.7±40.7cc, 671.5±79.8cc and 253.9±31.9cc, respectively. WMHv within each of the bi-lateral ACA, MCA and PCA territories were calculated for each subject and normalized by the subject’s total WMHv, resulting in a relative WMHv_rel_ for each territory. Figure 3 shows WMHv_rel_ for each vascular territory. Additionally, we combined both ACA and MCA to represent the combined anterior (ANT) territory. Out of the remaining 865 subjects, 162 did not have a stroke subtype classification available and were subsequently excluded from analysis.

**Figure 3:**
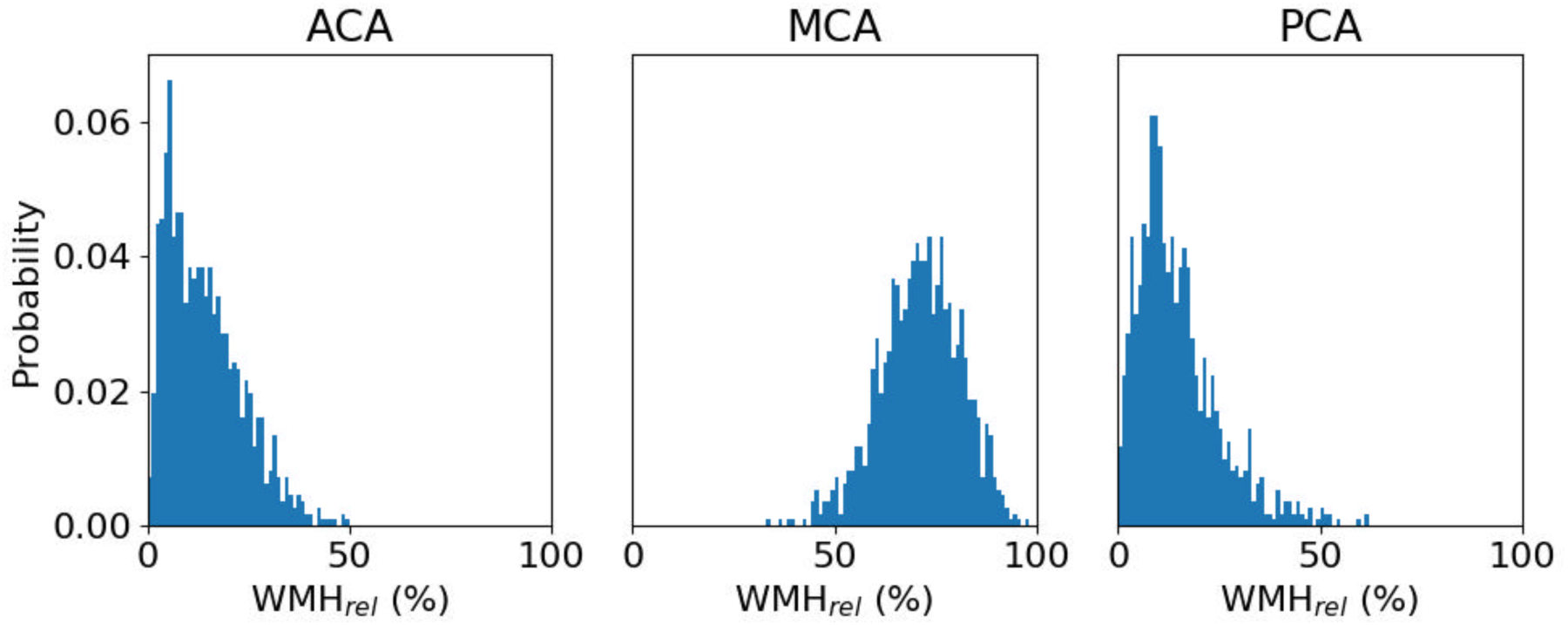
Distribution of relative WMH burden (WMHrel (%)) for each vascular territory, with a median WMHrel (interquartile range) of 13.1% (6.6%-20.7%), 70.7% (63.9%-77.5%) and 12.9% (7.8%-19.8%), respectively.

#### Clinical determinants of territorial WMHv: univariate analysis

Univariate analysis showed that age and territorial volume were significant correlates of relative WMH burden for all territories (p<0.001), while only HTN and sex (male) were found significant for ACA, PCA, and ANT territories (p≤0.001). Table 2 includes the complete set of univariate analyses.

**Table 2:**
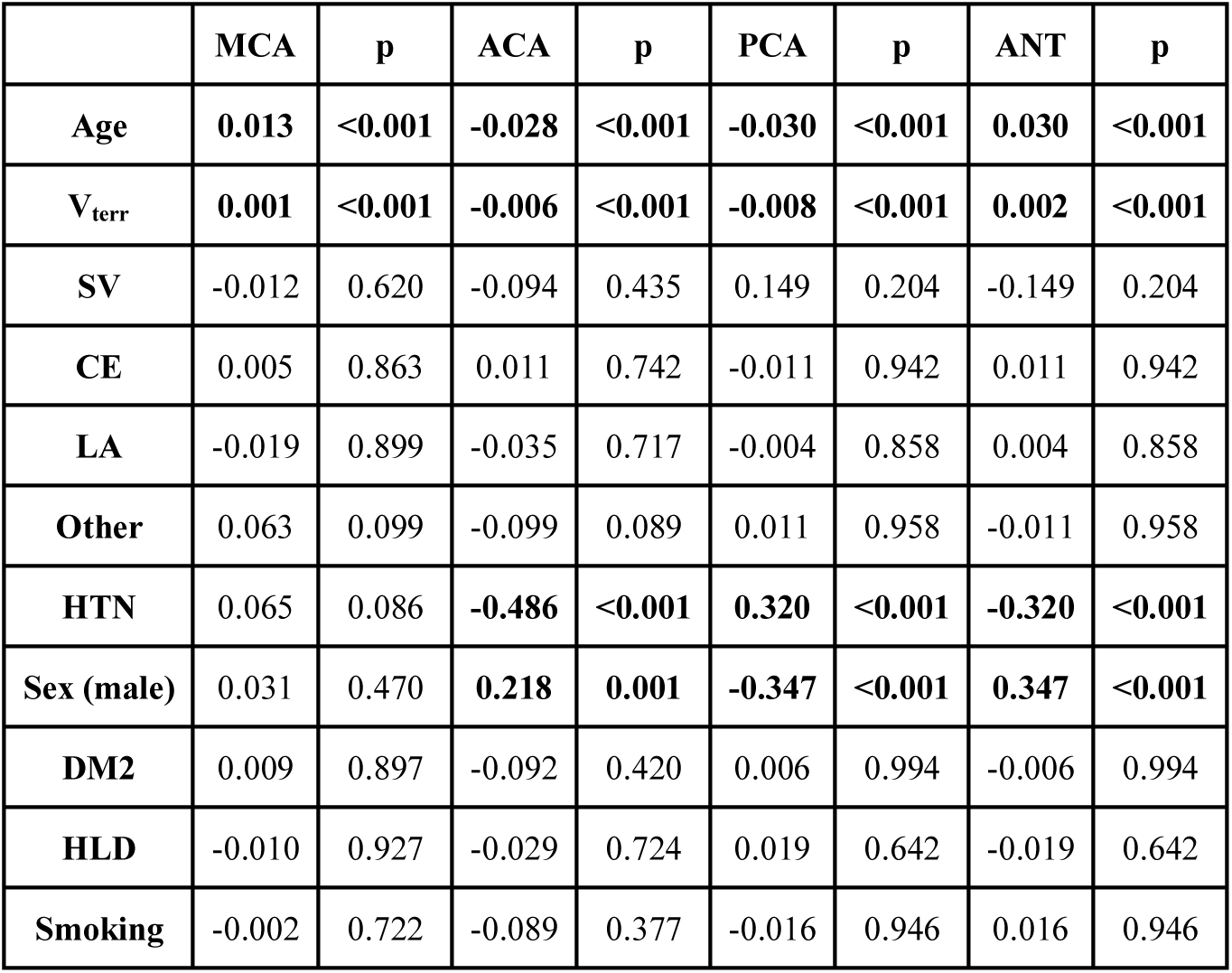
Univariate analysis of phenotypes with respect to relative WMH burden for each territory. Continuous variables were assessed using a linear model (reported beta and p-value), whereas categorical variables (maximum of two categories per variable) were assessed using a Mann-Whitney U Test (reported differences of the mean and p-value). Statistically significant correlations (p<0.05) are shown in bold.

|  | MCA | p | ACA | p | PCA | p | ANT | p |
| --- | --- | --- | --- | --- | --- | --- | --- | --- |
| Age | **0.013** | **<0.001** | **-0.028** | **<0.001** | **-0.030** | **<0.001** | **0.030** | **<0.001** |
| $V_{terr}$ | **0.001** | **<0.001** | **-0.006** | **<0.001** | **-0.008** | **<0.001** | **0.002** | **<0.001** |
| SV | -0.012 | 0.620 | -0.094 | 0.435 | 0.149 | 0.204 | -0.149 | 0.204 |
| CE | 0.005 | 0.863 | 0.011 | 0.742 | -0.011 | 0.942 | 0.011 | 0.942 |
| LA | -0.019 | 0.899 | -0.035 | 0.717 | -0.004 | 0.858 | 0.004 | 0.858 |
| Other | 0.063 | 0.099 | -0.099 | 0.089 | 0.011 | 0.958 | -0.011 | 0.958 |
| HTN | 0.065 | 0.086 | **-0.486** | **<0.001** | **0.320** | **<0.001** | **-0.320** | **<0.001** |
| Sex (male) | 0.031 | 0.470 | **0.218** | **0.001** | **-0.347** | **<0.001** | **0.347** | **<0.001** |
| DM2 | 0.009 | 0.897 | -0.092 | 0.420 | 0.006 | 0.994 | -0.006 | 0.994 |
| HLD | -0.010 | 0.927 | -0.029 | 0.724 | 0.019 | 0.642 | -0.019 | 0.642 |
| Smoking | -0.002 | 0.722 | -0.089 | 0.377 | -0.016 | 0.946 | 0.016 | 0.946 |

#### Clinical determinants of territorial WMHv: multivariate analysis

2/18/19 12:26:00 PMWe used the relative WMH burden as dependent variable and estimated the model parameters of the linear model, given by equation (1). This led to the removal (p-values for MCA, ACA, PCA, and ANT, respectively) of territorial volume (p = 0.615, 0.813, 0.583, 0.974), CE (p = 0.565, 0.506, 0.982, 0.982), LA (p = 0.817, 0.813, 1.000, 1.000), Other (p = 0.418, 0.787, 0.497, 0.497), and diabetes (p = 0.708, 0.199, 0.877, 0.877), refining the model to

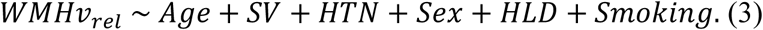

The results are summarized in Table 3 and diagnostic regression plots are shown in Figure 4. These results show that each phenotype modulates individual aspects of the spatial patterns of WMH burden in our cohort. In the ACA territory, age, SV stroke subtype, hypertensive status, HLD and smoking were found statistically significant, where HLD decrease the relative burden in ACA. The only phenotype found statistically significant in MCA is age, where an increase in age is associated with the relatively lower WMH burden in MCA. All phenotypes, except for hypertensive status, HLD, and smoking, were found statistically significant in PCA, as well as ANT.

**Figure 4:**
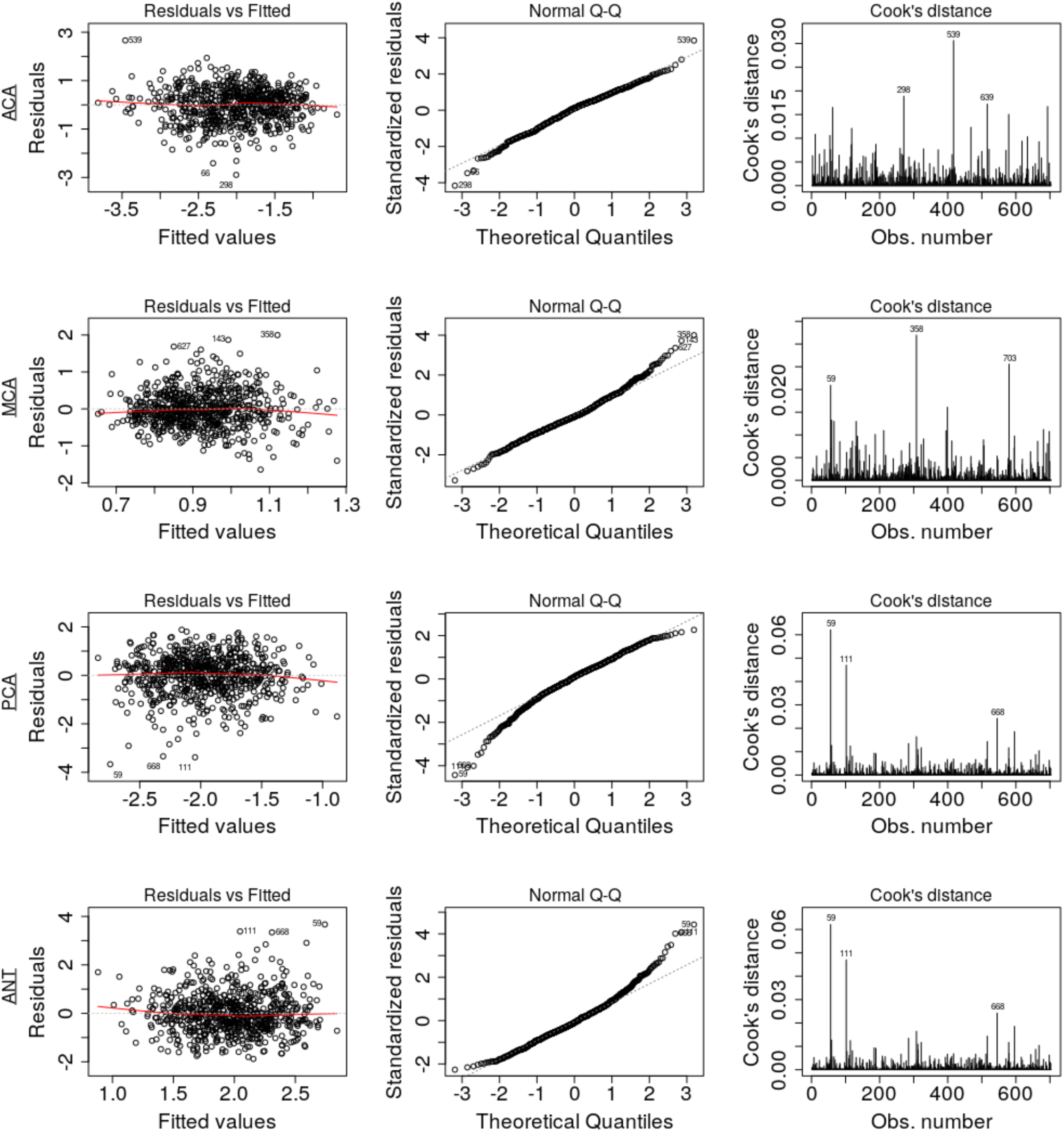
Diagnostic regression plots, resulting from the linear models for each of the four territories.

**Table 3:**
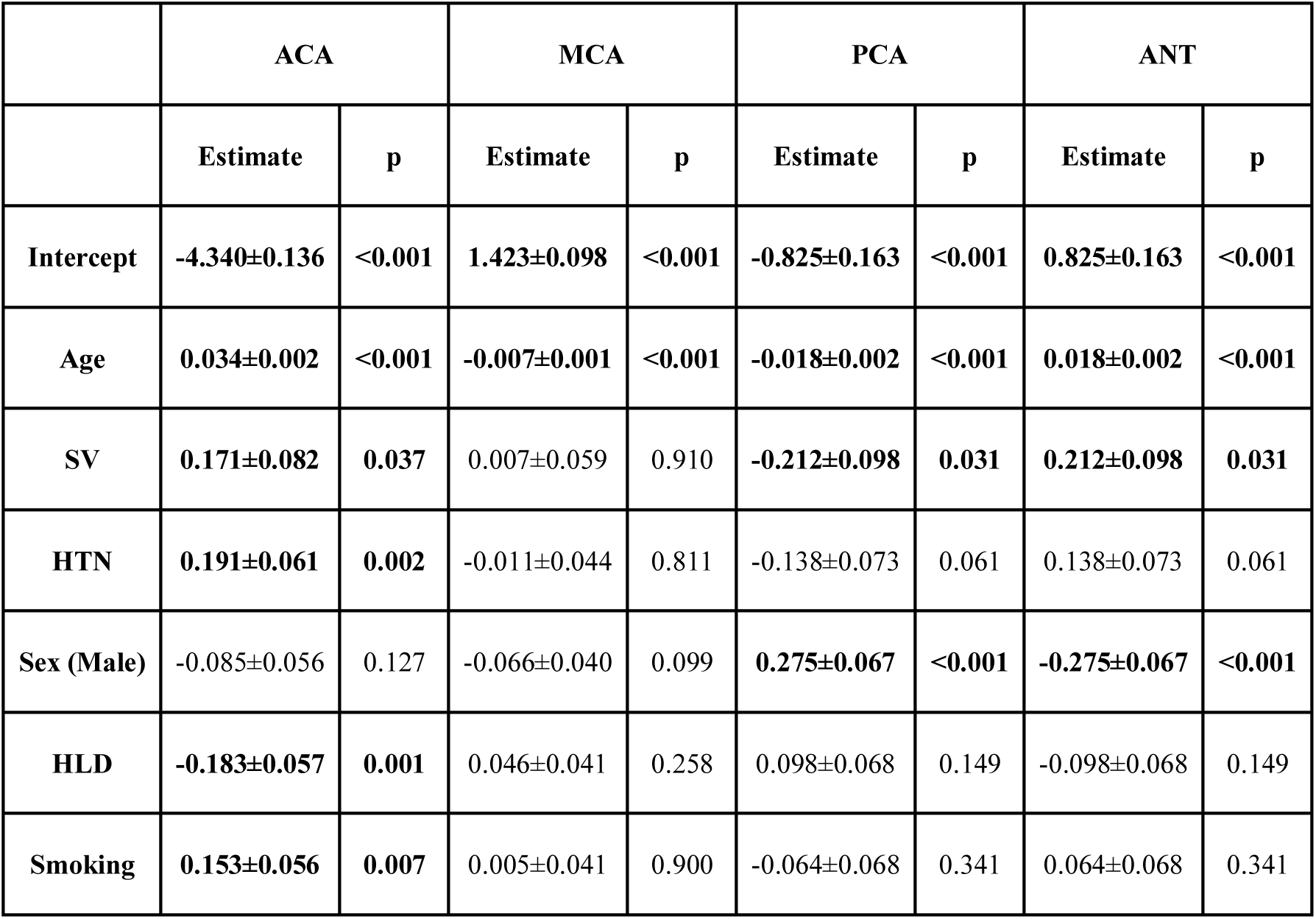
Parameter estimate for the linear model fit using R. Statistically significant parameters (p<0.05) are highlighted.

|  | ACA |  | MCA |  | PCA |  | ANT |  |
| --- | --- | --- | --- | --- | --- | --- | --- | --- |
|  | Estimate | p | Estimate | p | Estimate | p | Estimate | p |
| Intercept | -4.340±0.136 | <0.001 | 1.423±0.098 | <0.001 | -0.825±0.163 | <0.001 | 0.825±0.163 | <0.001 |
| Age | 0.034±0.002 | <0.001 | -0.007±0.001 | <0.001 | -0.018±0.002 | <0.001 | 0.018±0.002 | <0.001 |
| SV | 0.171±0.082 | 0.037 | 0.007±0.059 | 0.910 | -0.212±0.098 | 0.031 | 0.212±0.098 | 0.031 |
| HTN | 0.191±0.061 | 0.002 | -0.011±0.044 | 0.811 | -0.138±0.073 | 0.061 | 0.138±0.073 | 0.061 |
| Sex (Male) | -0.085±0.056 | 0.127 | -0.066±0.040 | 0.099 | 0.275±0.067 | <0.001 | -0.275±0.067 | <0.001 |
| HLD | -0.183±0.057 | 0.001 | 0.046±0.041 | 0.258 | 0.098±0.068 | 0.149 | -0.098±0.068 | 0.149 |
| Smoking | 0.153±0.056 | 0.007 | 0.005±0.041 | 0.900 | -0.064±0.068 | 0.341 | 0.064±0.068 | 0.341 |

Assumptions of the linear models, i.e. mean residual equals 0, no correlation between residuals and dependent variables, positive variability, homoscedasticity, and no multicollinearity (defined as variance inflation factors < 2), were fulfilled. However, autocorrelation of the residuals was significant for MCA, PCA, and ANT models. By removing the influential data points for the model fit using Cook’s distance for outlier detection (threshold 4/703), we identified 81 subjects (see Table 4) significantly younger and with larger WMH burden in the MCA territory. All assumptions for the linear models were fulfilled after removal of these subjects. Table 5 summarizes the difference in model fit before and after exclusion of the 81 subjects, showing a loss of significance in the stroke subtype for all models, and reveals significance of sex (male) in the ACA territory, as well as hypertensive status and hyperlipidemia in the PCA and ANT territories.

**Table 4:**
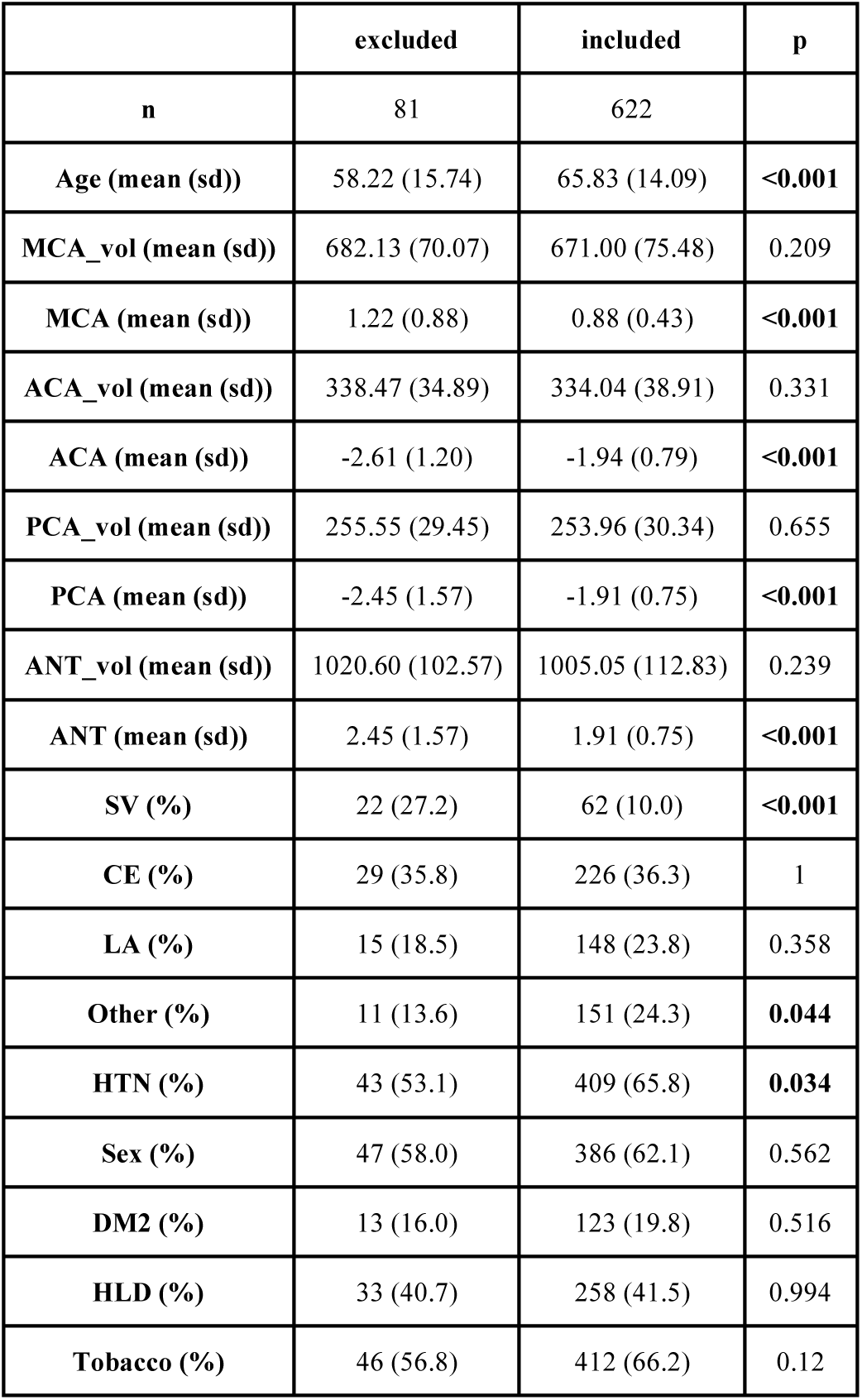
Characteristics of excluded subjects based on Cook’s distance threshold (4/703), compared to the remainder (included).

|  | excluded | included | p |
| --- | --- | --- | --- |
| **n** | 81 | 622 |  |
| **Age (mean (sd))** | 58.22 (15.74) | 65.83 (14.09) | **<0.001** |
| **MCA_vol (mean (sd))** | 682.13 (70.07) | 671.00 (75.48) | 0.209 |
| **MCA (mean (sd))** | 1.22 (0.88) | 0.88 (0.43) | **<0.001** |
| **ACA_vol (mean (sd))** | 338.47 (34.89) | 334.04 (38.91) | 0.331 |
| **ACA (mean (sd))** | -2.61 (1.20) | -1.94 (0.79) | **<0.001** |
| **PCA_vol (mean (sd))** | 255.55 (29.45) | 253.96 (30.34) | 0.655 |
| **PCA (mean (sd))** | -2.45 (1.57) | -1.91 (0.75) | **<0.001** |
| **ANT_vol (mean (sd))** | 1020.60 (102.57) | 1005.05 (112.83) | 0.239 |
| **ANT (mean (sd))** | 2.45 (1.57) | 1.91 (0.75) | **<0.001** |
| **SV (%)** | 22 (27.2) | 62 (10.0) | **<0.001** |
| **CE (%)** | 29 (35.8) | 226 (36.3) | 1 |
| **LA (%)** | 15 (18.5) | 148 (23.8) | 0.358 |
| **Other (%)** | 11 (13.6) | 151 (24.3) | **0.044** |
| **HTN (%)** | 43 (53.1) | 409 (65.8) | **0.034** |
| **Sex (%)** | 47 (58.0) | 386 (62.1) | 0.562 |
| **DM2 (%)** | 13 (16.0) | 123 (19.8) | 0.516 |
| **HLD (%)** | 33 (40.7) | 258 (41.5) | 0.994 |
| **Tobacco (%)** | 46 (56.8) | 412 (66.2) | 0.12 |

**Table 5:** Parameter comparison (significant) of all 4 models using the full data set and the subset, after 81 have been excluded based on Cook’s distance.

|  | ACA |  | MCA |  | PCA |  | ANT |  |
| --- | --- | --- | --- | --- | --- | --- | --- | --- |
|  | Full | Subset | Full | Subset | Full | Subset | Full | Subset |
| Intercept | -4.340<br>±0.136 | -4.330<br>±0.127 | 1.423<br>±0.098 | 1.443<br>±0.089 | -0.825<br>±0.163 | -0.765<br>±0.148 | 0.825<br>±0.163 | 0.765<br>±0.148 |
| Age | 0.034<br>±0.002 | 0.035<br>±0.002 | -0.007<br>±0.001 | -0.009<br>±0.001 | -0.018<br>±0.002 | -0.018<br>±0.002 | 0.018<br>±0.002 | 0.018<br>±0.002 |
| SV | 0.171<br>±0.082 | 0.116<br>±0.079 | - | - | -0.212<br>±0.098 | -0.090<br>±0.093 | 0.212<br>±0.098 | 0.090<br>±0.093 |
| HTN | 0.191<br>±0.061 | 0.181<br>±0.055 | - | - | -0.138<br>±0.073 | -0.168<br>±0.064 | 0.138<br>±0.073 | 0.168<br>±0.064 |
| Sex (Male) | -0.085<br>±0.056 | -0.134<br>±0.050 | - | - | 0.275<br>±0.067 | 0.226<br>±0.059 | -0.275<br>±0.067 | -0.226<br>±0.059 |
| HLD | -0.183<br>±0.057 | -0.163<br>±0.050 | - | - | 0.098<br>±0.068 | 0.124<br>±0.059 | -0.098<br>±0.068 | -0.124<br>±0.059 |
| Smoking | 0.153<br>±0.056 | 0.159<br>±0.051 | - | - | - | - | - | - |

Comparing the models summarized in Table 5 to their corresponding baseline models given by equation (2), we find that the reduction of residual deviation of the observed data is statistically significant in each case (p<0.001).

## 4 Discussion

In this work, we investigated the relative WMH disease burden per established cerebral vascular territory in AIS patients and how it is affected by common vascular risk factors, including age and sex. To allow automated characterization of WMH burden per vascular territory, we introduced an age-appropriate template with a 3D-FLAIR sequence for registration. Both WMH and ventricular volume of the template were representative of the disease burden in our AIS cohort and ventricular size of that found in healthy elderly (Ambarki et al., 2010). Moreover, this template allowed us to delineate vascular territories. We used this territorial map to elucidate relative spatial WMHv distributions in 865 patients with clinical 3D-FLAIR sequences, while accounting for size differences in the vascular territories.

Spatially specific effects of each investigated phenotype were identified in our cohort. Age shows a relative increase in the ACA and ANT with a relative decrease in the MCA and PCA territories. Age is known to be one of the most robust predictors of the overall WMH burden (Zhang et al., 2015). Our data suggest furthermore that an effect between the anterior and posterior vascular territory exists, where older patients accumulate more WMH burden in the ACA territory. A similar, but greater effect can be observed in case of patients with small-vessel stroke subtypes. While the exact biology of the underlying disease processes that drive these differences is unclear, spatial pathology distribution differences with posterior predominance were noted in patients with genetically defined small-vessel disease phenotypes such as CAA (Grabowski et al., 2001; Smith et al., 2008), which manifest early as impaired vascular reactivity and later as WMH and cerebral microbleeds. In these patient populations enriched for specific genetic mutations, small vessel dysfunction occurs early in life and appears to affect posterior circulation vessels first. Anterior circulation preponderance of the WMH burden seen in our study may imply that, by the time these patients present with a symptomatic event, their sporadic cerebral microangiopathy is diffuse and without predilection for posterior territory.

Smoking status shows a modifying effect on WMH burden with a relative increase in the ACA territory alone. Ghatan et al. (Ghatan et al., 1998) observed a reduced regional cerebral blood flow in the anterior cingulate cortex, among other regions. While there are no definite explanations as to the cause of the increased WMH burden yet, our findings, combined with those of Ghatan et al. may suggest chronic ischemia related to smoking leading to leukoaraiosis (O’sullivan et al., 2002). Sex (male) demonstrated the opposite effect, where larger relative burden were found in the posterior territory, similar but to a lesser extent than that of hyperlipidemia. There are no known biological mechanisms yet to explain these spatial differences.

In addition to smoking, hypertensive status shows an increase of relative WMH burden in the ACA territory, while a corresponding increase in ANT and decrease in the PCA territory only appeared after the subset analysis. There was no effect on the MCA territory, implying that the effect is driven by the spatial distribution of WMH in the ACA territory predominantly. The general lack of significance in modifying risk factors in MCA may be the result of the stroke lesions being predominantly in the MCA territory, thereby introducing uncertainty in identifying underlying disease burden (see Figure 2). The difference between ACA and PCA, however, may be due to differences between the small vessel architecture in both territories. Detailed investigations into those differences are urgently needed.

There are several limitations to our study. In general, spatial normalization of clinical low-resolution images is a non-trivial task, in particular in AIS patients, where biological responses to stroke, such as mass effects, can play an important role. In this study, we manually assessed the registration, leading to 17 subjects being excluded from final analysis due to gross registration errors. Nonetheless, small registration errors may be important for assessing WMHv within vascular territories. However, using an age-appropriate FLAIR template is a first step to mitigate the resulting uncertainties. Furthermore, using the template developed from MRI scans of age- and sex-matched individuals, who are stroke-free but with known intrinsic cerebrovascular pathology (CAA), has its advantage due its similarity to the average stroke patient brain. Although we demonstrated spatially discriminative changes of relative WMH burden, the underlying etiology needs to be further elucidated. In fact, accounting for common vascular risk factors that contribute to various ischemic stroke subtypes as well as adjusting for these TOAST subtypes further narrowed down pathophysiological contributions to WMH burden accumulation in the anterior versus posterior circulation. By removing influential data points before model fit we demonstrated the consistency of the presented trends. The majority of model parameters before and after excluding the 81 subjects from analysis fall within one standard deviation of another. However, small vessel occlusion become non-significant, whereas sex (ACA model), HTN (PCA and ANT model), and HLD (PCA and ANT model) become significant. While removing influential data points before model fit can bias the data, our results suggest that the risk factors with changing levels of significance should be interpreted with care. An additional limitation to our study was the lack of phenotype availability as related to other characteristics of small vessel disease such as chronic lacunes, cerebral micorbleeds and dilated perivascular spaces. Future use of these phenotypes may further characterize underlying brain pathology that contributes to variability in WMH accumulation by vascular territories and other anatomical distributions. While beyond the scope of this work, additional investigation into the capillary densities and networks of each vascular territory would also further this line of research. Another promising direction to delineate underlying WMH pathologies is to assess the effect of territorial mapping to border zone (“watershed”) areas between ACA-MCA and MCA-PCA territories. Given the hypothesis of diminished cerebral blood flow contribution to pathophysiology of WMH, a future study that develops and validates a concept of border zone through pathological-radiographic correlation would provide further insights on the role of chronic hypoxia-hypoperfusion etiology of WMH.

Other types of analyses will also be considered in future work. One promising, hypothesis free approach is the use of voxel-based lesion symptom mapping. While such analysis may reveal interesting correlates of the presentation of WMH in our cohort, its hypothesis free nature makes it difficult to directly interpret the results. By relying on a hypothesis-driven approach in this analysis, however, we are able to attribute the WMH burden to the individual supplying arteries.

The strengths of this analysis include (a) development of a novel FLAIR-based vascular territory template for clinical image registration in patients with stroke; (b) utilization of a large, thoroughly ascertained hospital-based cohort of AIS patients with detailed neuroimaging analysis; and (c) use of the validated methodologies for image processing and analysis.

In this work, we demonstrated that vascular risk factors influence spatial specificity of WMH, one of the most important radiographic manifestations of chronic cerebral ischemia. Here, we illustrated that WMH burden does not develop homogeneously throughout the supratentorial brain. These findings of spatial specificity of WMH in relation to vascular territory and risk factor exposure in AIS patients open the path for new investigations into underlying pathology of this common vascular disease and its associated risk factors, and ultimately into its connection with stroke.

## Acknowledgements

This project has received funding from the European Union’s Horizon 2020 research and innovation programme under the Marie Sklodowska-Curie grant agreement No 753896 (M.D. Schirmer). This study was supported by the NIH–National Institute of Neurological Disorders and Stroke (K23NS064052, R01NS082285, and R01NS086905), American Heart Association/Bugher Foundation Centers for Stroke Prevention Research, and Deane Institute for Integrative Study of Atrial Fibrillation and Stroke.

